# Stable isotopes and nanoSIMS single-cell imaging reveals soil plastisphere colonizers able to assimilate sulfamethoxazole

**DOI:** 10.1101/2024.01.11.575183

**Authors:** Qian Xiang, Hryhoriy Stryhanyuk, Matthias Schmidt, Hans H. Richnow, Yong-Guan Zhu, Li Cui, Niculina Musat

## Abstract

The presence and accumulation of both plastics and antibiotics in soils may lead to the colonization, selection and propagation of bacteria with certain metabolic traits e.g. antibiotic resistance, in plastisphere. However, the impact of plastic-antibiotic tandem on the soil ecosystem functioning, particularly on microbial function and metabolism remains currently unexplored. Herein, we investigated the competence of soil bacteria to colonize plastics and to mineralize/degrade ^13^C-labelled sulfamethoxazole (SMX). Using single cell imaging, isotope tracers, soil respiration and SMX mineralization bulk measurements we show that microbial colonization of polystyrene (PE) and polyethylene (PS) surfaces takes place within the first 30 days of incubation. Morphologically diverse, microorganisms were colonizing both plastic types, with a preference for PE substrate. Nano-scale Secondary Ion Mass Spectrometry measurements show that ^13^C enrichment was highest at 130 days with values up to 1.29 atom %, similar to those of the ^13^CO_2_ pool (up to 1.26 atom%, or 22.55 ‰). Our results provide direct evidence demonstrating, at single cell level, the capacity of bacterial colonizers of plastics to assimilate ^13^C from ^13^C-SMX. These findings expand our knowledge on the role of plastisphere microbiota in the ecological functioning of soils impacted by anthropogenic stressors.

## 1. Introduction

Global plastic pollution is a major challenge facing humankind as plastic is ubiquitously and persistently present in all environmental compartments (Horejs 2020). Unlike well-studied marine and fresh water environments, plastics in soil was only recently subject to investigations although it represents a significant amount, approximately 14% of the global plastic pollution (Wanner 2021). Plastic is expected to enter soil ecosystems through the plastic mulching, landfill, diffuse littering, and application of sewage sludge (Rillig 2012, Chae and An 2018, Zhang et al. 2019). Persistent in terrestrial ecosystems, plastic can accumulate and affect soil properties e.g. the soil bulk density, porosity, hydraulic conductivity, field capacity and plant performance as well as microbial community composition and fertility (Chae and An 2018, Agathokleous et al. 2021, Liu et al. 2021, Li et al. 2022). Abundant and distinct bacterial communities were shown to colonize plastic debris in soil i.e soil plastisphere (Zhang et al. 2019). These communities were distinct from those in the surrounding environment evidencing plastic debris as a selective habitat for microbial colonization in farmland soil (Zhang et al. 2019). Moreover, microbial community composition of the soil plastisphere via the high-throughput sequencing showed a high taxonomic diversity of the plastisphere colonizers (Bandopadhyay et al. 2020, Zhu et al. 2021, Xiang et al. 2022). With the potential to be transported together with the plastic debris over a broader spatial area, plastisphere microbial colonizers represent an emerging perturbation to complex environmental habitat, with yet unknown biogeochemical and ecological consequences. For instance, increasing evidence show that microbes living on the plastic surfaces have the potential to degrade the plastics and plastic additives, and may carry antibiotic resistance genes and human pathogens, impacting the ecosystem biochemistry and functioning (Zumstein et al. 2018, Rogers et al. 2020, Yang et al. 2020, Li et al. 2021, Zhu et al. 2021). Yet, the function and metabolic potential of microbial colonizers of soil plastisphere remain enigmatic. Although plastics are considered chemically inert, they can readily adsorb co-existing organic and inorganic pollutants such as antibiotics (Xiang et al. 2019).

The widespread use of antibiotics in humans and animals has significantly promoted the accumulation of antibiotics in a variety of environments. Antibiotic residues typically exert no significant acute toxicity in the environment, but induce the evolution and selection of antibiotic resistance genes within microorganisms, which pose a great threat to human health (D’Costa et al. 2011, Bottery et al. 2021, Murray et al. 2022). Recent studies suggested that plastics represent an increasing anthropogenic surface, which provide an avenue of enriching both microbes and antibiotics (Zettler et al. 2013, Wright et al. 2020, Zhu et al. 2021). Moreover, when the adsorbed antibiotics get in contact with the plastisphere microbiota, such chemicals may play important roles in restructuring the microbiota and therefore their ecological functions. A previous review indicated that the biofilm forming bacteria are protected against the bactericide effects of antibiotics, suggesting that antibiotics could induce specific biofilm formation, which have a defensive reaction (Stewart 2002), thus making plastisphere a potential selective habitat for antibiotic degraders. In addition, a previous study reported that higher microbial biomass and enzyme activity and a lower affinity for the substrate were found in the plastisphere compared to those of the rhizosphere, which indicated a stronger and faster carbon and nutrient turnover in the soil plastisphere (Zhou et al., 2021). However, no data is currently available whether plastic colonizers play a vital role in the transformation of co-occurring pollutants e.g. antibiotics.

The combination of fluorescent *in situ* hybridization (FISH) approaches, stable isotope probing and nano-scale Secondary Ion Mass Spectrometry (SIP-nanoSIMS) provides direct evidence for simultaneous detection of *in situ* phylogenetic identity, metabolic activity and function in complex microbial communities, at single cell level, without the need of cultivation (Fike et al. 2008, Li et al. 2008, Musat et al. 2008, Musat et al. 2012). In the present study, we conducted microcosm experiments using polyethylene (PE) and polystyrene (PS) plastic debris which were exposed over a time course of 130 days to soil amended with ^13^C-labelled SMX. Using Scanning Electron Microscopy (SEM) and Catalyzed Reporter Deposition-Fluorescence in situ Hybridization (CARD-FISH) with domain specific probes, we aimed to analyze morphology and abundances of microbial colonizers of plastic sheets with single cell resolution. Furthermore, we used SIP-FISH-nanoSIMS single-cell combinatory approach, to quantify the uptake of the ^13^C-labelled SMX by individual bacterial cells. The single-cell results combined with additional soil respiration and SMX mineralization bulk measurements provided novel evidence that bacterial colonizers of plastics are involved in the mineralization or partial transformation of the co-occurring antibiotics.

## 2. Materials and methods

### 2.1. Chemicals, plastic types and soil sampling

Sulfamethoxazole (IUPAC: 4-Amino-N-(5-methylisoxazol-3-yl)-benzenesulfonamide) was purchased from Sigma-Aldrich, USA. While SMX labeled with ^13^C at all six carbon atoms of the benzene ring (^13^C_6_-SMX) (Figure S1) was purchased from Clearsynth, India, with chemical purity of 97.69 % and isotopic enrichment of 99.13 %. All used solvents and chemicals, including hexamethyldisilazane (HMDS, Lot # STBJ2938), sodium cacodylate buffer (0.2M, pH = 7.4), were obtained from Merck in pro analysis quality.

Polyethylene (PE) and polystyrene (PS) are the two most commonly mass-produced polymers worldwide (PlasticsEurope, 2012), and widely used as mulch film in agriculture to enhance crop production by suppressing weeds, conserving soil water and increasing soil temperature. Here, commercial low-density PE (REWE, Germany) and PS (GoodFellow, England) films were selected as model plastics to conduct the incubation experiments. In preparation for the incubation experiments, the plastics were cut with sterilized scissors to produce 20 mm × 20 mm plastic sheets. Prior to the start of the experiments, optical profilometer was used to check the plastic surface roughness and select those with less than 10 µm roughness as suitable for microcosm incubations and further microscopy and spectrometry analyses. The information of the plastic surface roughness is shown in the supplementary information (Figure S2). Additionally, Raman spectroscopy was used to show that PE and PS have chemical-free surfaces (Figure S3), while the SEM indicated microbial-free plastic surfaces prior to the start of the experiment (Figure S4).

Surface soils (0−20 cm) were collected from an arable land in Xiamen, Fujian Province, China. After sampling, soil samples were immediately transferred to the laboratory and air-dried in a soil sample drying room at 20°C for several days prior to homogenisation. The air-dried soil was thoroughly mixed and sieved through a 2 mm mesh to remove plant debris and stones. After that, soils were moistened with sterile water to 60% of field capacity and pre-incubated at 25°C in the dark for 14 days to activate soil microorganisms before setting up the microcosm experiment.

### 2.2. SMX mineralization experiments and ^13^CO_2_ production

The concentration of SMX used in this work was 40 mg kg^-1^ soil which is double in comparison with SMX concentration used by previous study that successfully characterized the SMX-degraders in soil (Ouyang et al. 2019). For analyzing the mineralization of SMX in soil, a parallel batch experiment in closed soil microcosms with the variants (1) control soil (without acetone and SMX), (2) soil mixed with acetone, (3) soil amended either with ^12^C-SMX or ^13^C-SMX, were incubated for a period of 30 days. The mineralization experiments were conducted in 500 ml Schott flasks sealed with an OxiTop® - Respirometer (WTW, Germany) for determination of oxygen consumption. A NaOH solution was used to trap the formed CO_2_ (Figure S5). The isotope measurements and concentration analyses are described in the supporting information S1.

### 2.3. Laboratory microcosms & Experimental set-up

Based on the results of the tracer experiment for SMX mineralization, microcosms were established to allow plastic colonization by soil native microorganisms during an incubation experiment in soil (Figure S6): Soil controls were plastic sheets introduced in soil without SMX, Soil + ^12^C_SMX (plastic sheet introduced in soil amended with unlabeled SMX), Soil + ^13^C_SMX (plastic sheet introduced in soil amended with ^13^C labeled SMX). For the amendment we prepared SMX stock solution of 2 g L^−1^ with either unlabeled or ^13^C_6_-labeled SMX in acetone. Four ml of the stock solution was spread drop by drop to 10 g soil (give final concentration 800 mg kg^-1^ soil) and mixed thoroughly, followed by air drying for 2 h to evaporate the acetone. Finally, 10 g of the SMX-amended dried soil was well mixed with 190 g of activated soil (give final concentration 40 mg kg^-1^ soil). All treatments were conducted in 250 ml glass baker microcosms. Within each microcosm containing 200 g soil, 5 plastic sheets of 20 × 20 mm of each plastic type, PE and PS film respectively, were inserted. All the microcosms were kept at 28°C in dark and a humidity of 14.5% was maintained with sterile water over a total period of 130 days constantly. At selected time points during the incubation (30, 70, and 130 days), the plastic sheets were taken out from the soil with sterilized forceps and chemically fixed with 15 ml 2% (v/v) paraformaldehyde (PFA) (Sessitsch et al.) in 0.2 M cacodylate buffer (CB, pH 7.4) overnight at 4°C.

### 2.4. Scanning electron microscopy (SEM)

The PFA-fixed plastic pieces were washed 3 times with CB solution, followed by dehydration in an ethanol series in CB of 30, 50, 70, 80, 90, 96, and 100% (3min each). Subsequently, the plastic pieces were dried with 50% Hexamethyldisiloxane in ethanol and 100% Hexamethyldisiloxane solution respectively (10 min each). Prior to SEM imaging, the plastic sheets were sputtered with Au/Pd mixture and evaluated with a Scanning Electron Microscope (Merlin VP Compact, Carl Zeiss, Germany). Imaging was performed with a secondary electron detector at a working distance of 2.0 mm and an electron high tension of 2.0 kV.

### 2.5. Fluorescence in situ hybridization (FISH) coupled with catalyzed reporter deposition (CARD-FISH)

Following the PFA fixation, the plastic coupons were washed 3 times with CB solution, dehydrated in 30 and 50% ethanol in CB (3 min each) and further stored in 50% ethanol in CB at 4°C until used for CARD-FISH procedure. The CARD-FISH was performed as described elsewhere (Musat et al. 2014). Briefly, plastics were coated with 0.2% low-melting point agarose to avoid cell loss by detaching from the plastic surface during the following steps of permeabilization, hybridization and washing. Plastisphere cells were permeabilized with lysozyme (10 mg ml^−1^in 0.05 M EDTA, pH 8.0; 0.1 M Tris–HCl, pH 7.5) for 1h at 37°C, followed by washing in sterilized ultrapure water and treated with 60 U/ml achromopeptidase 30 min at 37°C. Endogenous peroxidases were inactivated by incubation of plastics in 3% H_2_O_2_ in sterilized water for 10 min at room temperature (RT). The plastic pieces were hybridized with HRP-labeled EUB338 (Amann et al. 1990) probe for 3 h at 46°C in a hybridization buffer containing 0.9 M NaCl, 20 mM Tris–HCl (pH 7.5), 10% (w/v) dextran sulfate, 0.02% (w/v) SDS, 35% (v/v) formamide (Fluka) and 1% (w/v) blocking reagent (Boehringer, Mannheim Germany). The HRP-probe concentration was 0.166 ng ml^−1^ (probe stock solution of 50 ng ml^−1^diluted 1:300 v/v in hybridization buffer). Hybridized plastics were incubated for 15 min at 48°C in pre-warmed washing buffer. The CARD step was performed for 20 min at 46°C in the dark in standard amplification buffer containing 1 μg ml^−1^ Alexa Fluor 594-labelled tyramides (Thermo Fisher Scientific). Meanwhile, parallel plastic coupons were hybridized with nonsense probe NON EUB338, to account for false positive signals (Wallner et al. 1993). The hybridized plastic sheets were further stained for 10 min with 1 μg ml^−1^ of 4′, 6′-diamidino-2-phenylindol (DAPI). For fluorescence microscopy investigation, embedding in a 4:1 (v/v) mixture of low fluorescence glycerol mountant (Citifluor AF1, Citifluor) and mounting fluid VectaShield (Vecta Laboratories) was applied. Hybridizations were evaluated by fluorescence microscopy using an Axio Imager. Z2 microscope (Carl Zeiss) and filter sets for DAPI and Alexa Fluor 594 dyes.

### 2.6. Nano-scale Secondary Ion Mass Spectrometry (nanoSIMS)

Given the presence of soil particles on the plastic surfaces that will render the cell identification difficult, the CARD-FISH fluorescence micrographs were used to define and map areas of interest for single-cell analysis by nanoSIMS. The nanoSIMS analysis was done to assess the metabolization of SMX by quantifying the incorporation of ^13^C by individual bacterial cells (identified as such by EUB probe hybridization) colonizing the plastic surfaces at 30, 70 and 130 days. For the nanoSIMS analysis, the hybridized and imaged plastics were directly mounted on the nanoSIMS sample holder and coated with 20 nm Au/Pd (80/20 %) layer to provide a conductive surface and then analyzed with a NanoSIMS-50L instrument (CAMECA, AMETEK) in negative extraction mode employing 16 keV Cs^+^ primary ions. Prior to the analysis, implantation of cesium was done via pre-sputtering of 120×120 μm sample area with 200 pA of 16 keV Cs^+^ ion beam for 16 min to enhance the yield of negative secondary ions. For the analysis, 2 pA Cs^+^ ion beam was rastered in a 512×512 pixels saw-tooth pattern over 40 × 40 μm of pre-sputtered sample area with 2 ms dwell time per pixel. The secondary ions were analyzed with double-focusing magnetic sector mass spectrometers for their mass-to-charge ratio (m/z) and 7 secondary ion species (^12^C^12^C^−^, ^12^C^13^C^−^, ^12^C^14^N^−^, ^13^C^14^N^−^, ^31^P^−^, ^32^S^−^ and ^31^P^16^O^2−^) were collected in parallel. For each field of view, 80 scans were corrected for lateral drift, accumulated and further processed with the Look@NanoSIMS software (Polerecky et al. 2012). Regions of interest (Musat et al. 2008, Musat et al. 2014) (RoIs) were manually defined based on ^12^C^14^N^−^ secondary ion distribution maps and identified and cross-checked for the topographical and morphological appearance of single cells using fluorescence microscopy and FISH. The cellular carbon isotope enrichment upon labelling and the natural ^13^C abundance (range from 1.004 to 1.121 atom%) were derived as the fraction of heavy ^13^C^14^N^−^ and light ^12^C^14^N^−^ molecular ion counts, i.e. ^13^C^14^N^−^/(^12^C^14^N^−^+^13^C^14^N^−^), from single-cell confined RoIs.

### 2.7. Isotope ratio measurement of CO_2_ and soil organic carbon

The sample preparation for concentration CO_2_ measurement and analysis of soil organic matter is described in the Supporting information S1. The carbon isotope composition of CO_2_ was determined with a GC-combustion isotope ratio mass spectrometer (GC-IRMS, Thermo Finnigan MAT 253 253, Bremen, Germany). The isotope composition of soil organic matter was determined with an Elemental Analyzer coupled to isotope ratio mass spectrometer (Thermo Fisher Scientific, Bremen, Germany) (Girardi et al. 2013) Isotope data were reported in delta notation as described in Tamisier and colleagues (Tamisier et al. 2022).

## 3. Results and discussion

### 3.1. SMX mineralization and soil respiration

The SMX mineralization and soil respiration activity were analyzed during the first 30 days of incubation using isotope tracers. After 30 days incubation, the results showed that the total organic carbon in SMX-free and SMX amended soil amounted to 14.45 ± 1.48 mg kg^-1^ and 16.76 ± 1.22 mg kg ^-1^, respectively (Figure 1A). The ^13^C-soil carbon was ranging between −26.66 ‰ and 37.14 ‰ in ^12^C_6_-SMX and ^13^C_6_-SMX treated soil, respectively (Figure 1B). The soil respiration activity indicated by formation of CO_2_ was constantly decreasing during the 30 days incubation (Figure 1C). This indicates that the amendment of SMX reduce soil respiration, most probably by inhibiting microbial activity. The isotope composition of the CO_2_ in the control without SMX amendment showed typical soil respiration isotope values of −26.1 ± 1.3‰. The ^13^C_6_-SMX-amended soil incubations showed an increase of the δ^13^C value of CO_2_ in the gas phase from −9.9 to 22.6‰ and demonstrated that ^13^C_6_-SMX mineralization occurred (Figure 1D). These results are in accordance with previous work that characterized SMX-degraders in pig manure applied soil (Ouyang et al. 2019), although we have used a higher amount of antibiotics during incubations (40 mg kg^-1^, double in comparison to Ouyang et al.,) Also, much older reports suggested that microorganisms were able to survive and remain metabolically active under high antibiotic concentration (Stewart. 2002) supporting further our findings. Herein, formed microbial biofilms on the PE and PS plastic debris were observed during our experiments, which may have a protective role against the bacteriostatic effect of such high SMX concentrations during the prolonged exposure of up to 130 days. The continuous increase of ^13^CO_2_ over time from ^13^C_6_-SMX measured here indicates that SMX was mineralized or partially metabolized by microorganisms, possibly antibiotic resistant and/or antibiotic tolerant. This hypothesis implies that the SMX-mineralizing microorganisms, particularly those colonizing the plastics, should became gradually enriched in ^13^C. This was analyzed by the combination of stable isotope tracers and FISH-nanoSIMS single cell imaging.

**Figure 1.**
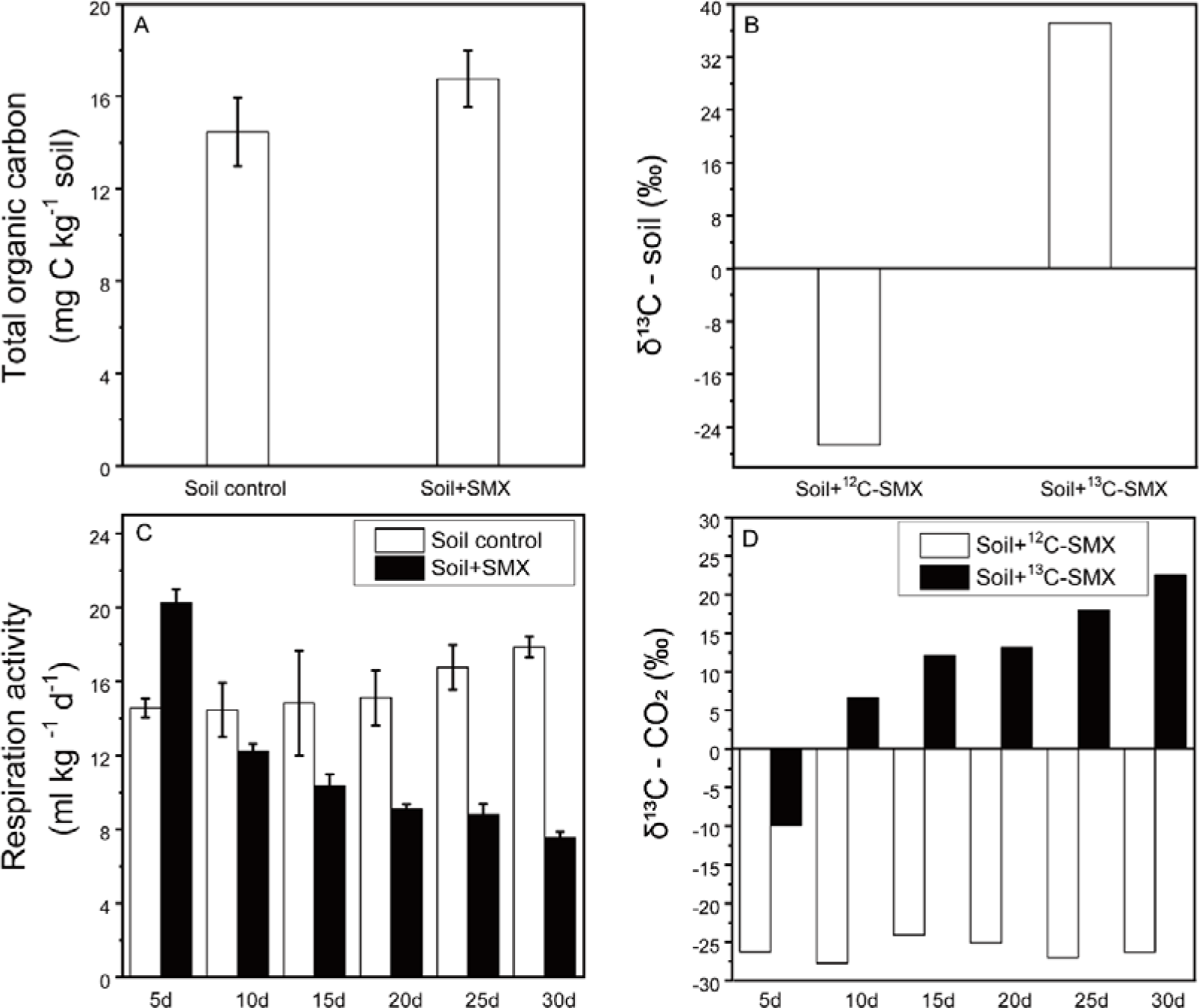
Sulfamethoxazole (SMX) mineralization and soil respiration. (A) The total organic carbon in soil; (B) δ^13^C value of soil carbon; (C) Effect of SMX on soil respiration activity; (D) Production of ^13^C-labeled CO_2_ over a period of 30 days, measured at 5-days intervals in soil microcosms amended with ^13^C-SMX or ^12^C-SMX.

### 3.2. Colonization of the plastisphere by the soil microorganisms

Prior to single-cell measurements by nanoSIMS, we applied SEM to assess and visualize microbial colonization and morphology of cells in the plastisphere (Figures S7-S11). We observed a high diversity of morphologically distinct microorganisms on the plastic coupons, including cocci-like, filamentous, and rod-shaped which were the three most prevalent morphotypes (Figures S7-S11). Moreover, we observed the prevalence of particular morphotypes preferentially colonizing PE or PS plastic foils with or without SMX amendment. Thus, PE plastic coupons without SMX load, visually showed abundant filament-like cells while predominantly cocci-like cells were observed on the PE plastic surfaces amended with SMX (Figure S11). In contrast, rod-like cells seem to be abundantly colonizing both types of plastic independent of SMX presence (Figure S11). Considering that our experiment was not designed for statistical analysis of the colonizing morphotypes, and only limited fields of view and plastic coupons were imaged, we can only report on the colonization trends observed during SEM imaging (S2 text). Of note, further studies need to consider the inherent colonization and distribution heterogeneity of microorganisms on such plastic surfaces when statistical analyses of cell morphology and distribution are planned. Based on SEM observations we conclude that the colonization of plastic sheets in the presence of antibiotics supports the hypothesis that such surfaces once released in the soil environment became attractive niches for rapid microbial colonization. In addition, the soil plastics may act as potentially selective surfaces for metabolically versatile microbial groups e.g. antibiotic resistant and/or tolerant bacteria such as those reported to be enriched in agricultural soils fertilized with animal manure. Previous SEM observations of plastic debris colonization in soil incubation experiments (Zumstein et al. 2018) and aquatic environments (Zettler et al. 2013, Harrison et al. 2014, Rogers et al. 2020) reported similar results of phenotypically diverse microorganisms comprising of fungi, cyanobacteria, heterotrophic bacteria etc. extensively colonizing plastic surfaces. Generally, the colonization and biofilm formation on plastic particles seems to be dictated by a variety of factors comprising environmental sample composition, substrate type, surface properties, sample location etc (Rogers et al. 2020). In our study, considering that initial chemical and microbial sample composition of the soil sample was the same in all incubation experiments, the plastic type and antibiotic presence/absence seems to be the key players influencing the colonization event by different morphotypes.

### 3.3. Cell abundance in the plastisphere

CARD-FISH using domain-specific Eubacterial probe was employed to determine total bacterial abundance in the PE- and PS-plastisphere. The results showed that the amendment of SMX significantly reduced the bacteria counts on both PE and PS plastisphere (ANOVA, *P* < 0.05). For example, in microcosm experiments without SMX addition, the density of bacteria on PE and PS surfaces ranged from 1.29E+10^4^ to 2.33E+10^4^, and 2.92E+10^3^ to 1.49E+10^4^ hybridized cells/mm^2^, respectively (Figure 2). While in the PE and PS plastisphere formed under SMX amendment, the hybridized bacterial cell counts were lower ranging from 5.36E+10^3^ to 2.06E+10^4^, and 2.06E+10^3^ to 3.43E+10^3^ hybridized cells/mm^2^, respectively (Figure 2). Additionally, by comparing the two types of plastic, we found that PE coupons harbored a higher number of hybridized bacterial cells than PS (ANOVA, *P* < 0.05) (Figure 2). Similar to the SEM observations, CARD-FISH findings suggest that bacteria prefer to colonize the PE coupon and to a lesser extent the PS, which indicate that plastic type may be one of the major factors driving the abundance and possibly phylogenetic and functional diversity of plastisphere in soil habitats. These findings are supported by previous observations of a higher abundance of ARGs reported in the soil PE-plastisphere compared to PS-plastisphere (Zhu et al. 2021), which may suggest the enrichment of SMX-resistant or tolerant microorganisms on PE surfaces. Taken together, our SEM and FISH imaging results suggest a plastic-type dependent colonization by soil microorganisms, with a preference for PE habitat colonization.

**Figure 2.**
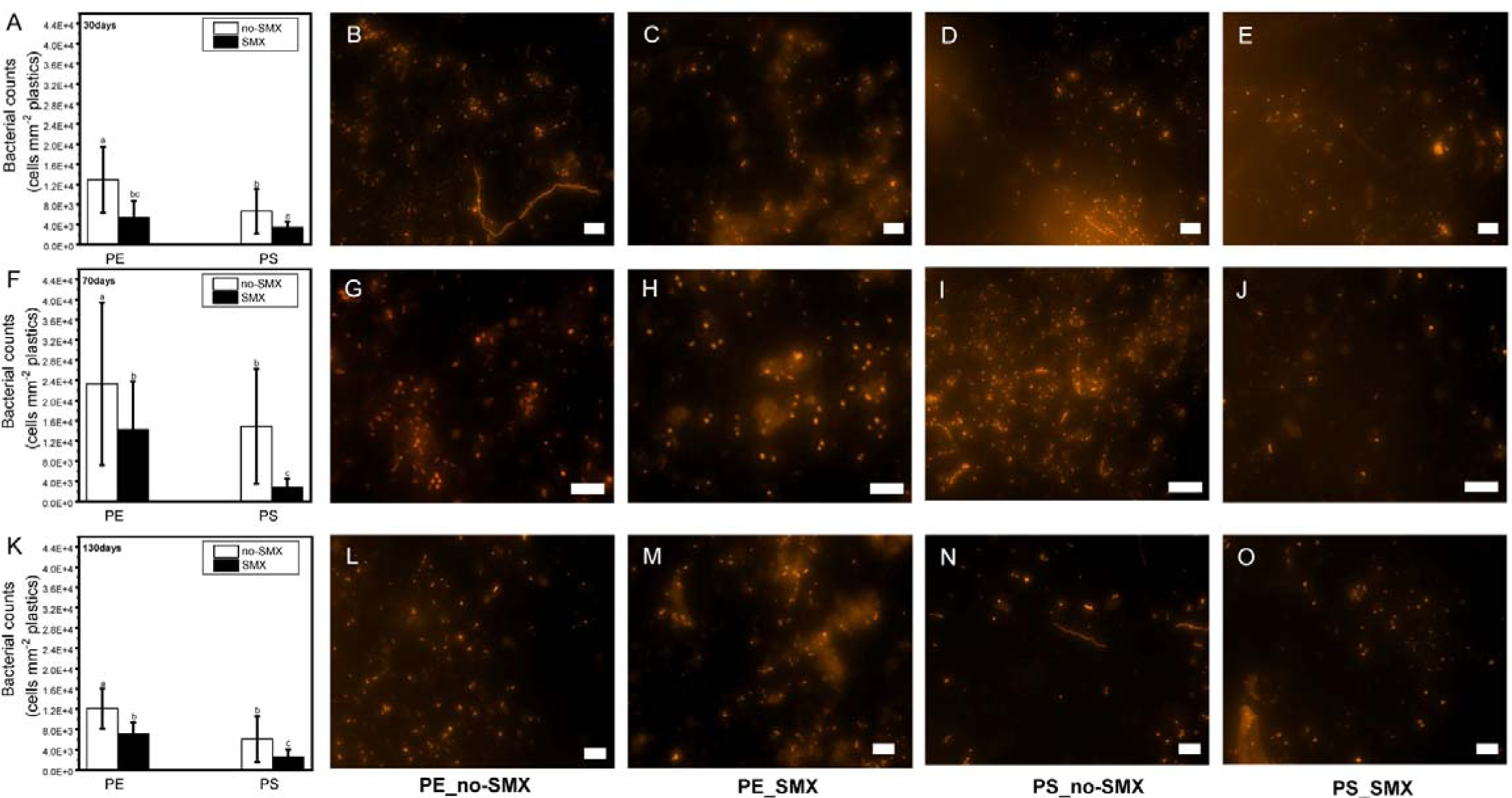
Cell abundance resulted from CARD-FISH with the HRP-labeled EUB338 probe and Alexa 594 tryamides. Bacterial abundances on PE and PS plastisphere over a period of (A) 30 days, (F) 70 days, and (K) 130 days incubation in soil microcosms without or with SMX addition. Representative epifluorescence micrographs of the microbial colonizers in PE plastisphere formed in soil microcosms without SMX addition over (B) 30 days, (G) 70 days, and (L) 130 days incubation and with SMX amended over (C) 30 days, (H) 70 days, and (M) 130 days incubation; Representative epifluorescence micrographs of the microbial colonizers in PS plastisphere without SMX amended over (D) 30 days, (I) 70 days, and (N) 130 days incubation and with SMX amended over (E) 30 days, (J) 70 days, and (O) 130 days incubation. Hybridized bacterial cells can be visualized as orange dots. The white scale bar represents 10 μm.

### 3.4. Single-cell NanoSIMS anaylsis

NanoSIMS analysis was conducted to i) determine if bacteria colonizing PE and PS plastisphere are capable of actively taking up ^13^C-SMX or ^13^C-SMX metabolites and to ii) quantitatively asses the ^13^C isotopic enrichment in individual cells and the number of ^13^C-enriched cells within the time frame of the experiment as direct prove of their involvement in SMX transformation/degradation. To ensure identification of bacterial cells for single-cell nanoSIMS measurements, CARD-FISH and fluorescence microscopy was applied prior to nanoSIMS (Figure 3). Single-cell ^13^C-fraction in ^13^C-SMX amended experiments was quantified relative to single-cell measurements of bacteria colonizing plastic surfaces in the absence of SMX. The ^13^C fraction (atom %) was derived from ^12^C^14^N/^13^C^14^N isotope ratio of single-cells nanoSIMS measurements. The ^13^C fraction of bacterial colonizers of plastic surfaces in the microcosm experiments without SMX exposure was ranging from 1.004 to 1.121 atom%, with a median value of 1.056 atom% (Figure S12). We further used this range as the baseline for quantifying the ^13^C enrichment in cells colonizing the PE and PS plastisphere (Figures 4 and 5). In the labeling experiments, colonizing cells at 30, 70 and 130 days became gradually and slightly enriched in ^13^C; however we measured similar median values, ranging from 1.11 to 1.15 atom% for both plastic types across the incubation range (Figures 4 and 5). Notably, the number of single cells showing ^13^C enrichment increased in time, particularly those colonizing the PS surface at 130 days. Almost all cells analyzed (n ≥ 100) at this time point were enriched in ^13^C above the maximum value of the control cells (Figure 5H, Table S1). Generally, the PS surface showed a higher number of ^13^C enriched cells, while the PE showed a more abundant bacterial colonization (Figures 2, 4 and 5) suggesting that the plastic type may select for SMX mineralizing/degrading bacteria. The single cell ^13^C enrichment was highest at 130 days with values up to 1.29 atom %, very similar to the measured ^13^C enrichment values of the CO_2_ pool (up to 1.26 atom%, or 22.6 ‰) (Figures1, 4 and 5, S1 text). The relatively low increment of ^13^C enrichment suggests that the colonizing cells assimilate only a minor fraction of SMX-derived carbon, being most probably reliant on other common carbon sources present in soil. In addition, the slow degradation of antibiotics may cause a higher antibiotic resistance enrichment in soil plastisphere. Our results indicate that bacterial communities colonizing the plastisphere have the capacity to, at least partly, degrade and assimilate SMX-derived carbon into their biomass. To explain the relatively low ^13^C enrichment we propose three working models: i) bacterial colonizers of PE and PS mineralize or partially degrade the SMX slowly and directly assimilate the ^13^C-SMX derived carbon; ii) primary SMX degraders release labelled secondary metabolites which are slowly assimilated by other members of the community; and iii) primary SMX degraders generate ^13^CO_2_ which is assimilated by autotrophic members of the community. Given the low isotopic enrichment of cells, and the similarity between the ^13^C enrichment of single cells and the CO_2_ pool, the latest two models seem the most likely ones. This is supported by previous nanoSIMS-based studies showing that direct assimilation of labeled substrates usually leads to significantly higher biomass enrichment (up to 6 atom%) during comparable incubation times (Rotaru et al. 2018, Zumstein et al. 2018). In contrast, label transfer via secondary metabolic products such as organic carbon or CO_2_ may lead to much lower enrichment into the assimilating cells e.g. heterotrophs or autotrophs (Alonso et al. 2012, Arandia-Gorostidi et al. 2017, Vidal et al. 2018). Further systematic investigations of the plastisphere in soils are needed to identify the phylotypes involved in SMX-transformation and assimilation. Considering the low single-cell uptake of ^13^C-SMX reported here, high resolution imaging approaches e.g. SIP-FISH-nanoSIMS seems the most suited methodology to investigate such assimilation/degradation processes, rather than conventional SIP approaches which require a higher level of isotope labelling into cellular components (DNA, RNA or proteins).

**Figure 3.**
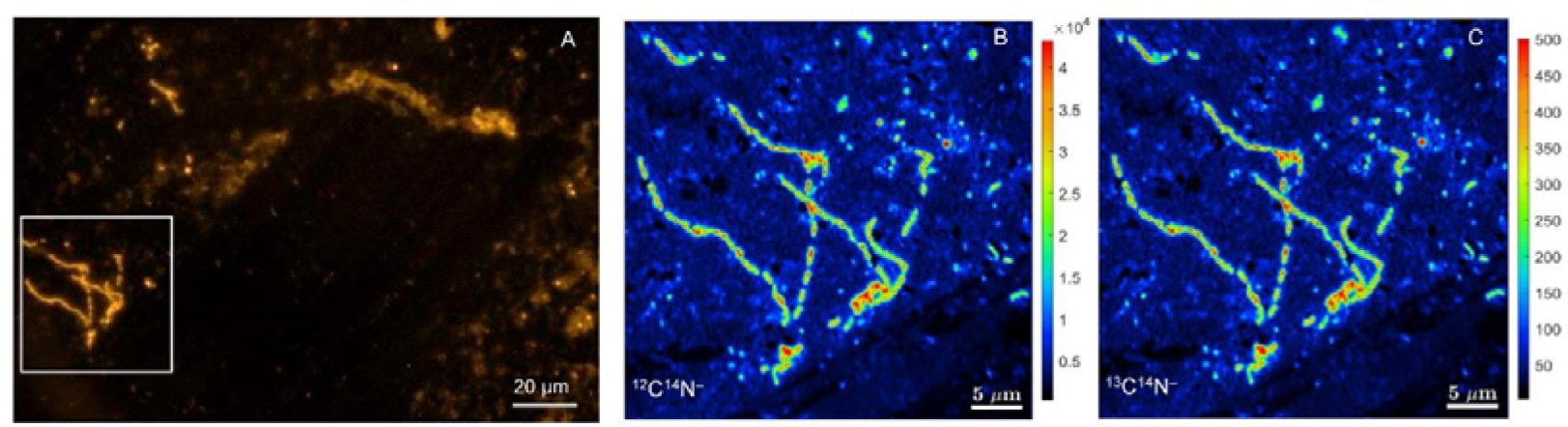
Correlative SIP-FISH-nanoSIMS imaging. (A) Representative epifluorescence micrograph of the microbial colonizers in plastisphere amended with ^13^C-SMX prior to CARD-FISH with the HRP-labeled EUB338 probe and Alexa 594 tryamides; (B), (C) NanoSIMS images of ^12^C^14^N^−^(B), and ^13^C^14^N^−^ (C) molecular ions show that cells in plastisphere were enriched in ^13^C.

**Figure 4.**
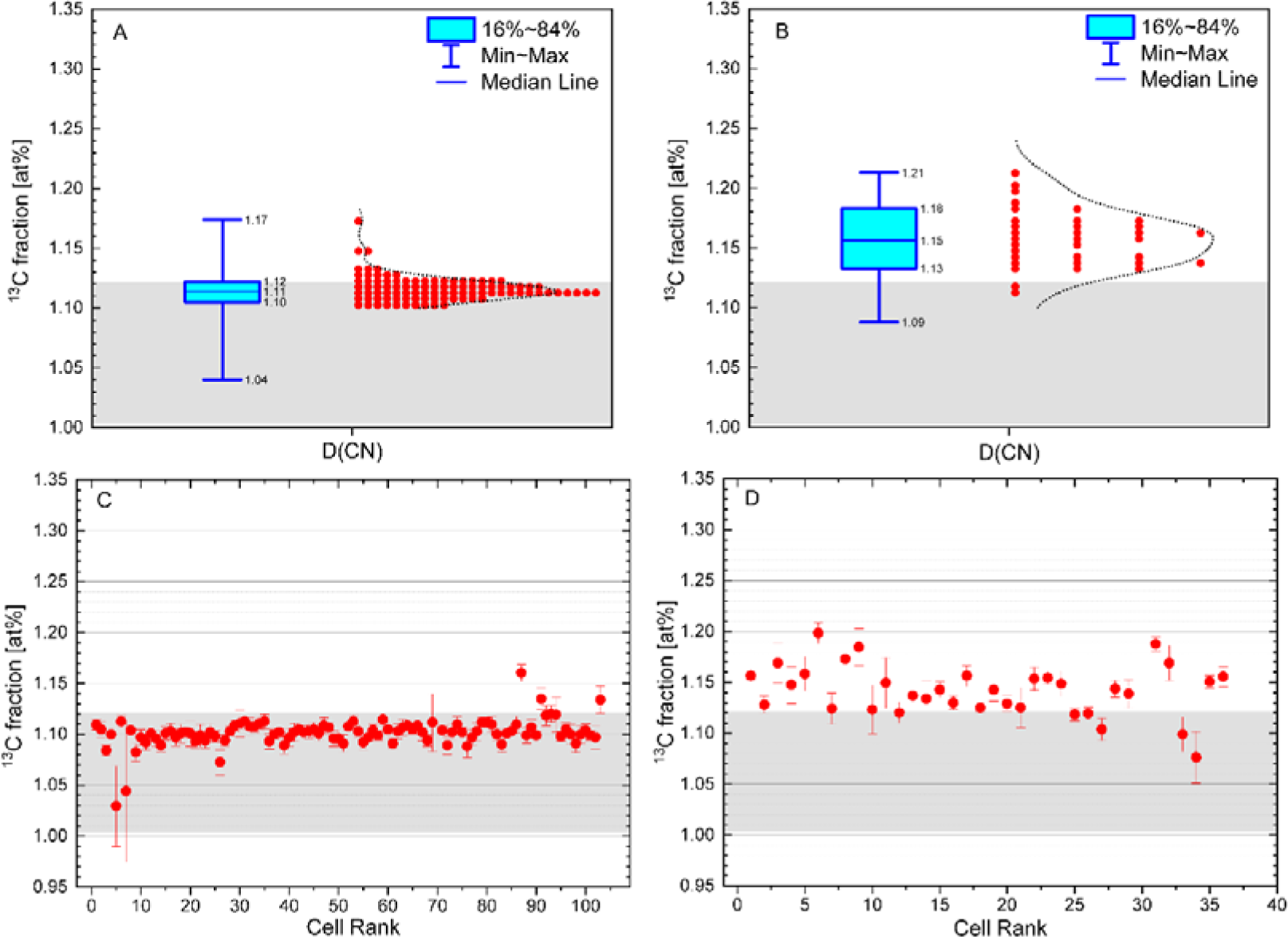
Distribution of single microbial cells based on the ^13^C fraction derived from ^12^C^14^N^−^ and ^13^C^14^N^−^ molecular ions measured by nanoSIMS on PE and PS plastic surfaces with 30 days incubation. The box plots showing the ^13^C fraction (atom%) of single cells in (A) PE and (B) PS plastisphere after 30 days incubation in soil microcosms. The scatter diagrams showing the ^13^C enrichment (mean ± sd) in the cellular groups in (C) PE and (D) PS plastisphere compared to the natural abundance of ^13^C fraction measured by NanoSIMS after 30 days incubation. The black grey rectangle represents the range of the natural abundance of ^13^C fraction from control measured cells (1.004 to 1.121 atom%). The box plots show the range of 16%-84% percentile (lower and upper box boudaries), the median value (line within the box), and the data minimum and maximum (whiskers). Each dot is a measurement of a single cell.

**Figure 5.**
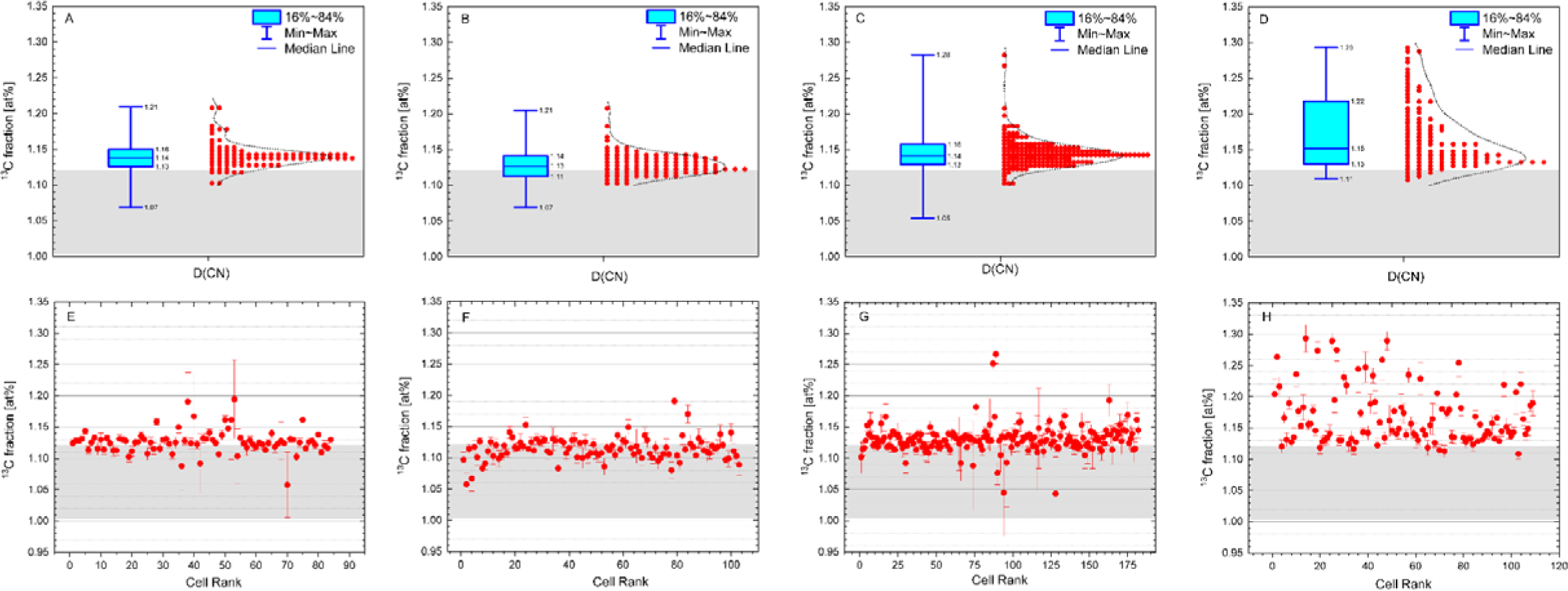
NanoSIMS quantitation showing the ^13^C fraction derived from ^12^C^14^N^−^ and ^13^C^14^N^−^ molecular ions of single microbial cells colonizing the PE and PS plastic surfaces during microcosms experiments with ^13^C_6_-SMX addition. The box plots showing the ^13^C fraction (atom%) of single cells measured by NanoSIMS in (A) PE plastisphere after 70 days, (B) PE plastisphere after 130 days, (C) PS plastisphere after 70 days, (D) PS plastisphere after 130 days incubation in the soil microcosms. The scatter diagrams show the ^13^C enrichment of single cells compared to the natural abundance of ^13^C fraction in (E) PE plastisphere after 70 days, (F) PE plastisphere after 130 days, (G) PS plastisphere after 70 days, (H) PS plastisphere after 130 days incubation in the soil microcosms. The box plots show the range of 16%-84% percentile (lower and upper box boudaries), the median value (line within the box), and the data minimum and maximum (whiskers). The dots were presented as the mean ± sd. Each red point is a measurement of a single cell. The black grey rectangle represents the range of the natural abundance of ^13^C fraction (1.004 to 1.121 atom%) measured in single cells on plastic surfaces in ^12^C_6_-SMX amended microcosm experiments.

Taken together, this study presents an experimental combinatory approach based on isotope tracers to study the metabolic capability of soil bacteria colonizing plastic surfaces to degrade sulfamethoxazole, a bacteriostatic antibiotic. The use of ^13^C labeled SMX was central to determine the role of plastisphere bacteria in SMX degradation by tracing SMX-derived carbon into both CO_2_ pool and bacterial biomass. Our results have implications in microbial and soil ecology and biotechnology. From an ecological perspective, we show that emerging contaminants like antibiotics and plastics cannot be transformed or completely biodegraded by single organisms, but rather by metabolic networking of colonizing organisms. Studying their intertwined function in a spatial context is key to understanding and further harvest their metabolic potential for novel biotechnology applications. Stable isotope tracers and chemical imaging approaches with single-cell and isotopic resolution such as nanoSIMS and Raman technology can guide the mining of function-targeted microorganisms colonizing plastics directly from the environment. Our study offers a beneficial edge of the plastisphere through its potential to select microbiomes that can be further harnessed to lower pollutants in natural and man-made habitats.

## 4. Conclusions

In the present study, we used stable isotope tracers coupled to single-cell imaging by nanoSIM, FISH and SEM to determine if natural microbial communities are able to colonize plastic surfaces in the presence of antibiotics and if these colonizers can degrade antibiotics. Our results shows that a morphologically rich microbial community is able to colonize plastic surfaces in the presence of antibiotics and can assimilate ^13^C-labelled SMX into cell biomass, the direct experimental proof of their active role in the degradation of plastic’ adsorbed antibiotics. Our findings bring new evidence on the functional and ecological role of plastisphere microbiota and their potential impact on the functioning of soil ecosystem and the governing microbial processes.

## Supporting information

supplemental information for the main text.

## Acknowledgment

This research was funded by National Natural Science Foundation of China (22193061, 42207143). We acknowledge the Centre for Chemical Microscopy (ProVIS) at the Helmholtz Centre for Environmental Research for using their analytical facilities. ProVIS is supported by European Regional Development Funds (EFRE – Europe funds Saxony). Further financial support was provided by the Chinese Scholarship Council.

