## supplemental information for the main text. for "Stable isotopes and nanoSIMS single-cell imaging reveals soil plastisphere colonizers able to assimilate sulfamethoxazole"

**S1 text**

**Mineralization of sulfamethoxazole (SMX) in soil**

Mineralization experiment was performed in 500 ml Schott bottles with three replicate bottles without SMX, two replicate bottles with $^{12}C$-SMX and one bottle with $^{13}C$-SMX. All the microcosms were incubated in the dark at 28 °C for 30 days. The formed $CO_2$ in the gas phase leaving the bottles was trapped in 20 ml 1 M NaOH solution. At 5, 15, 20, 25 and 30d, the NaOH was sampled and distributed to 20 ml serum bottles and the bottles were closed with Teflon-lined rubber stoppers and aluminum crimp caps for the determination of the total $CO_2$ (soil respiration) and its isotopic composition. Notably, before the measurement, the trapped $CO_2$ was totally released with excessed 6M HCl solution at pH >3. Then the concentration of the total $CO_2$ and the $^{13}C$ isotope composition of the $CO_2$ was determined by gas chromatography isotope ratio mass spectrometry (GC-IRMS, Thermo Finnigan MAT 253 253, Bremen, Germany), as described elsewhere [1]. After each NaOH collection, the bottles were flushed with air to ensure aerobic conditions and replaced with new NaOH solution. The average soil respiratory activity was calculated as $CO_2$ production per kilogram dry soil per day. The conversion of delta $^{13}C$ value (delta per mill) into $^{13}C$ fraction D [atom%] were calculated as follows: D=1/(1+1/[(1+0.001×delta)×0.01074529]) ×100 [atom%], where 0.01074529 is the reference value used for delta $^{13}C$ calculation. To validate the results of the GC-IRMS method, the total organic carbon and $^{13}C$ isotope compositions of the soils were determined by EuroEA3000 elemental analyzer (EuroVector, Milan, Italy) coupled with a MAT 253 isotope ratio mass spectrometer (Thermo Fisher Scientific,

Bremen, Germany).[2]

**S2 text**

**Colonization trends observed during SEM imaging**

Total density of cell-like features on PE and PS surfaces in the soil plastisphere with and without SMX addition. In the plastisphere from SMX addition experiments total density of cell-like features show and apparent increase 2 to 13 times, respectively from day 30, when colonization was first imaged, to day 130, when the experiment was stopped. Additionally, when comparing the two different types of plastic with SMX addition, it appears that PE film harbored a slightly higher number of cell-like features than PS.

**Fig. S1**

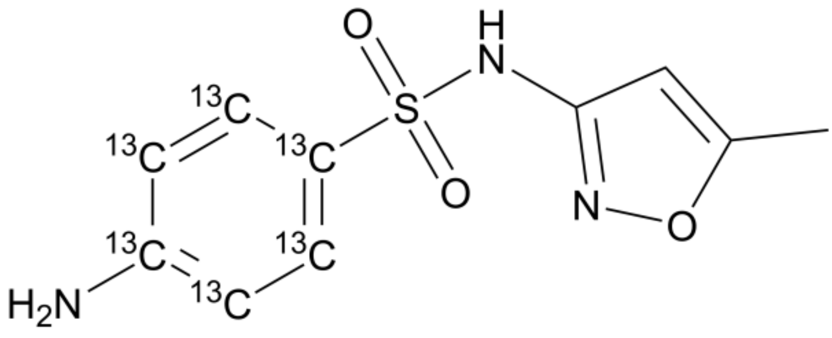


**Figure S1** The structure of $^{13}C_6$-SMX.

**Fig. S2**

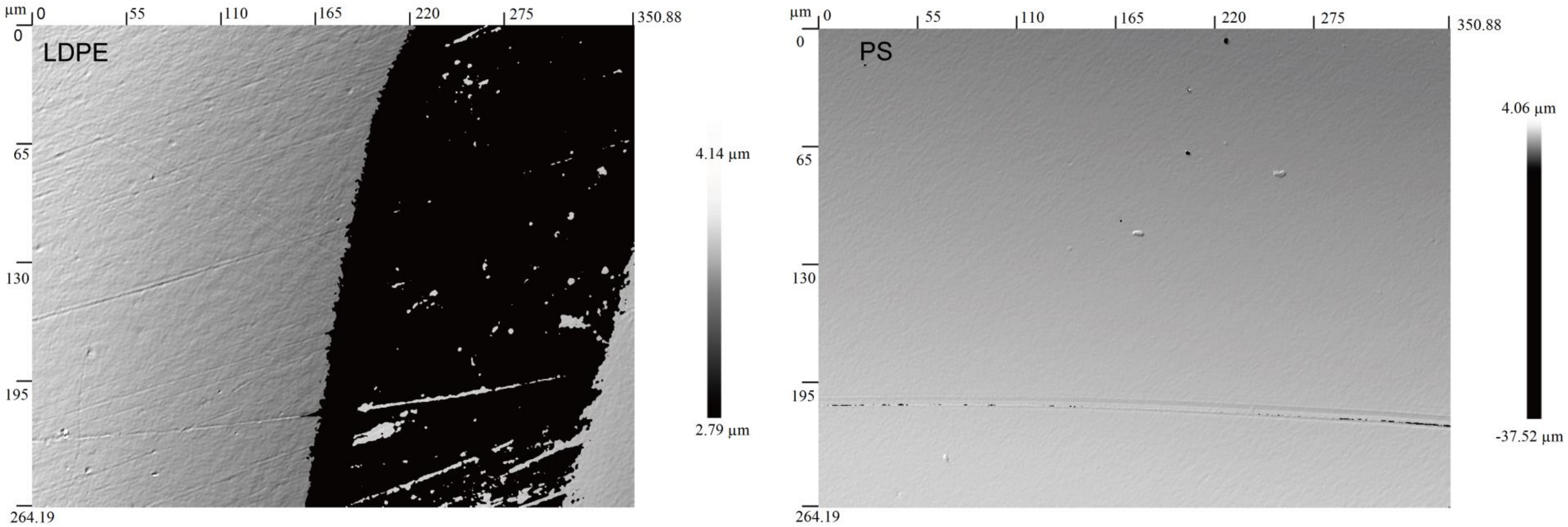


**Figure S2** Roughness of the low density polyethylene (LDPE) and polystyrene (PS) surfaces measured by optical profilometer.

**Fig. S3**

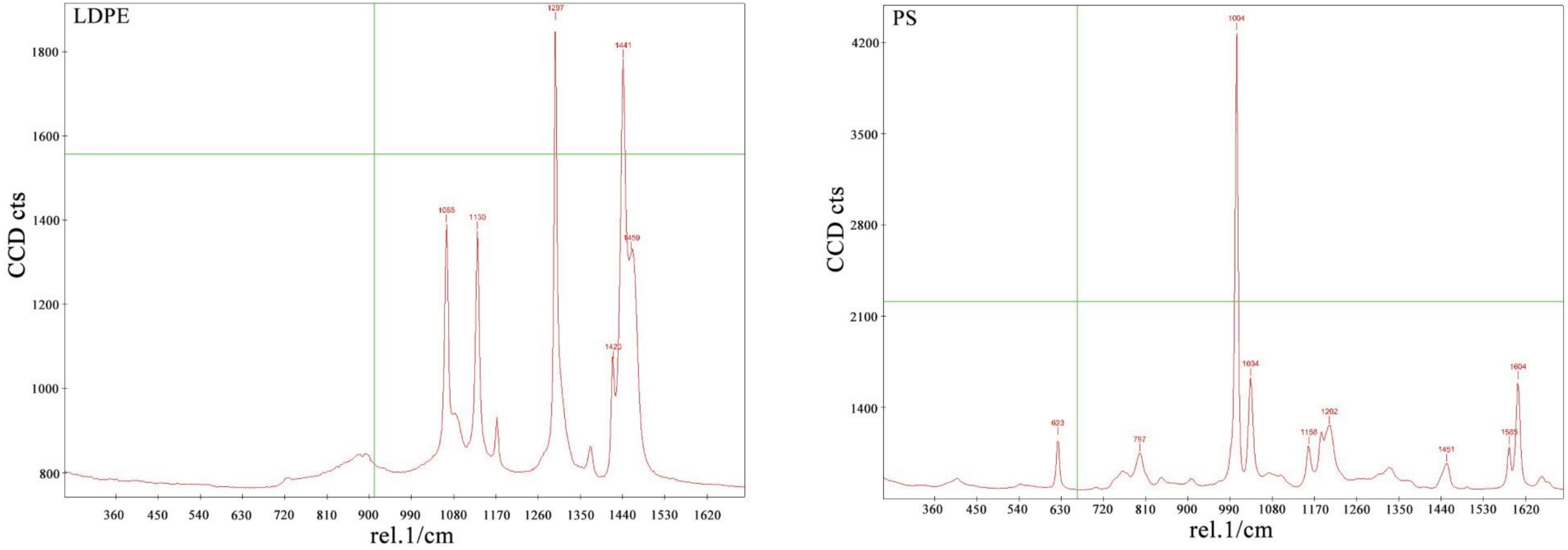


**Figure S3** Raman spectrum of the naked LDPE and PS plastic surface.

**Fig. S4**

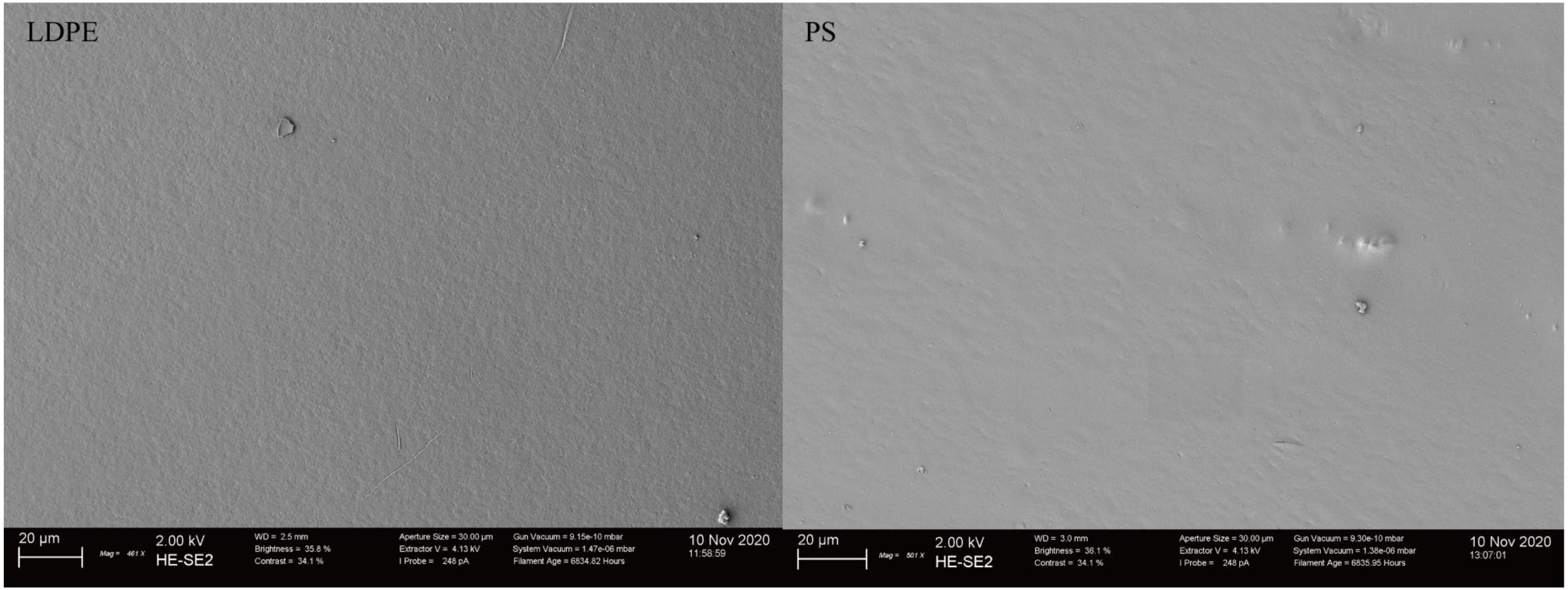


**Figure S4** SEM images showing the naked LDPE and PS plastic surface without microbes.

**Fig. S5**

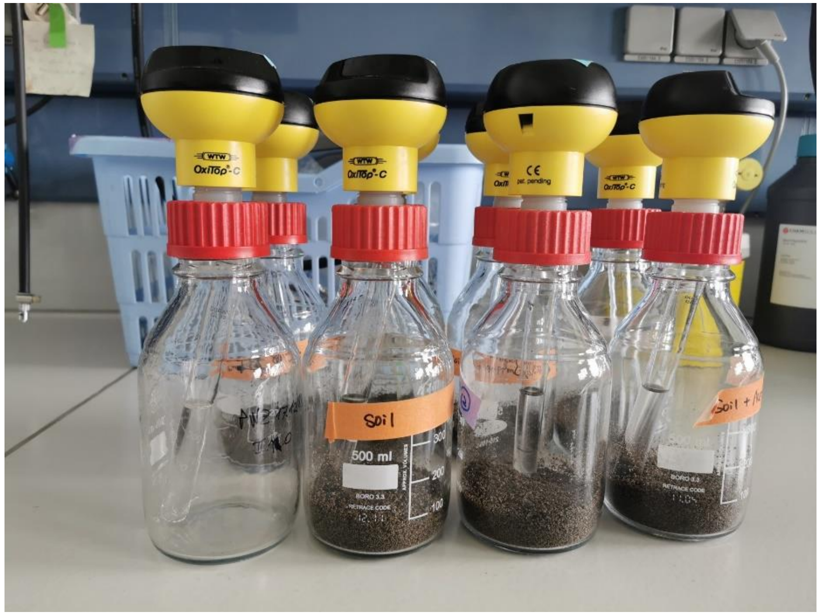


**Figure S5**. SMX mineralization in soil. Oxitop sensor for determination of oxygen consumption. NaOH traps for trapping $CO_2$ to analyze $CO_2$ production and SMX mineralization. The $CO_2$ concentration and $^{13}CO_2$ composition were subsequently analyzed by IR-MS.

**Fig. S6**

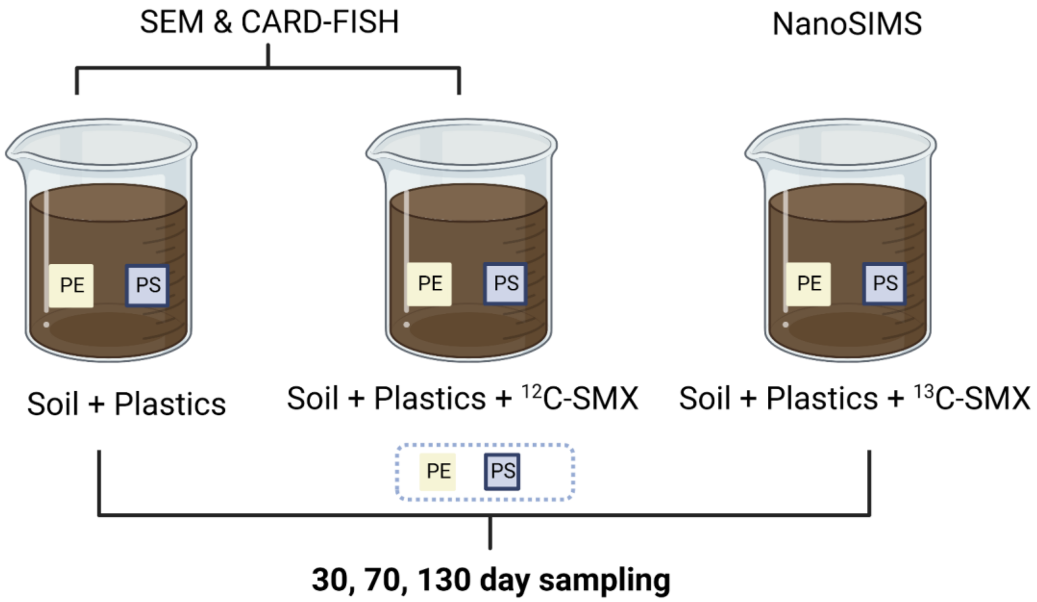


**Figure S6**. Experimental design and approaches used.

**Fig.S7**

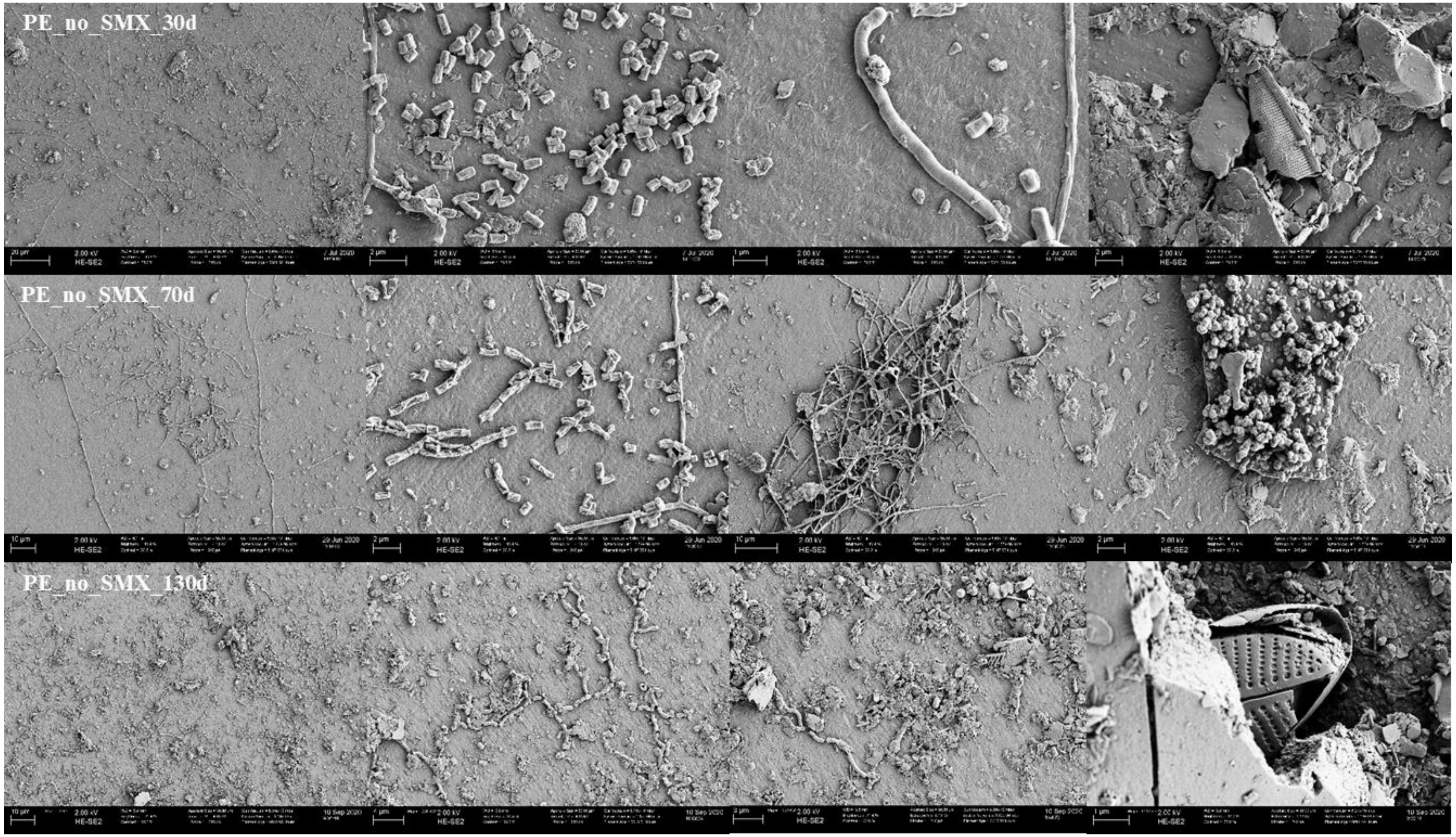


**Figure S7** SEM images of microbial colonization of PE substrate incubated in soil for 30, 70 and 130 days without SMX.

Fig. S8

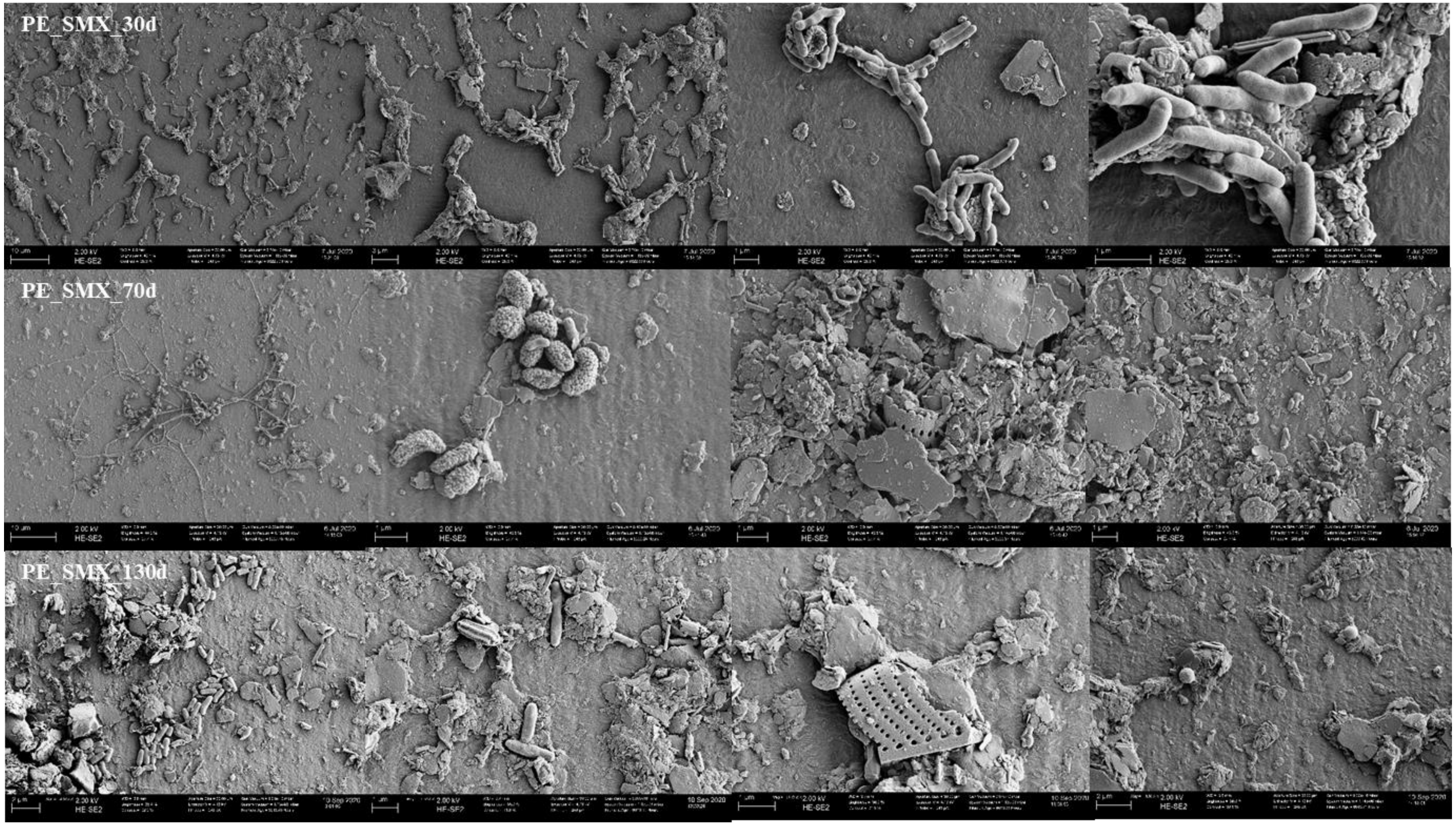


**Figure S8** SEM images of microbial colonization of PE substrate incubated in soil for 30, 70 and 130 days with SMX amendment.

**Fig.S9**

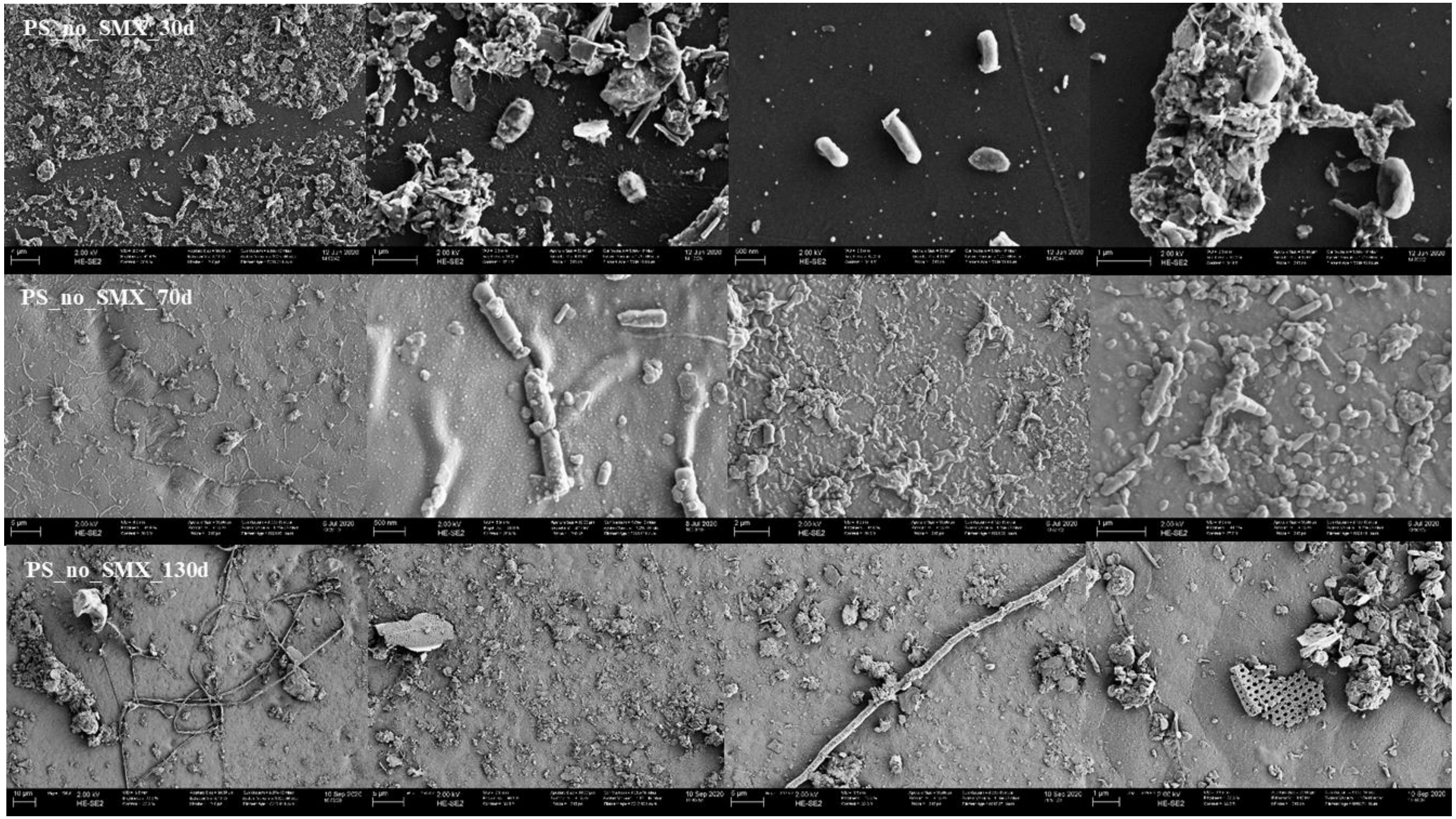


**Figure 9**. SEM images of microbial colonization of PS substrate incubated in soil for 30, 70 and 130 days without SMX.

**Fig. S10**

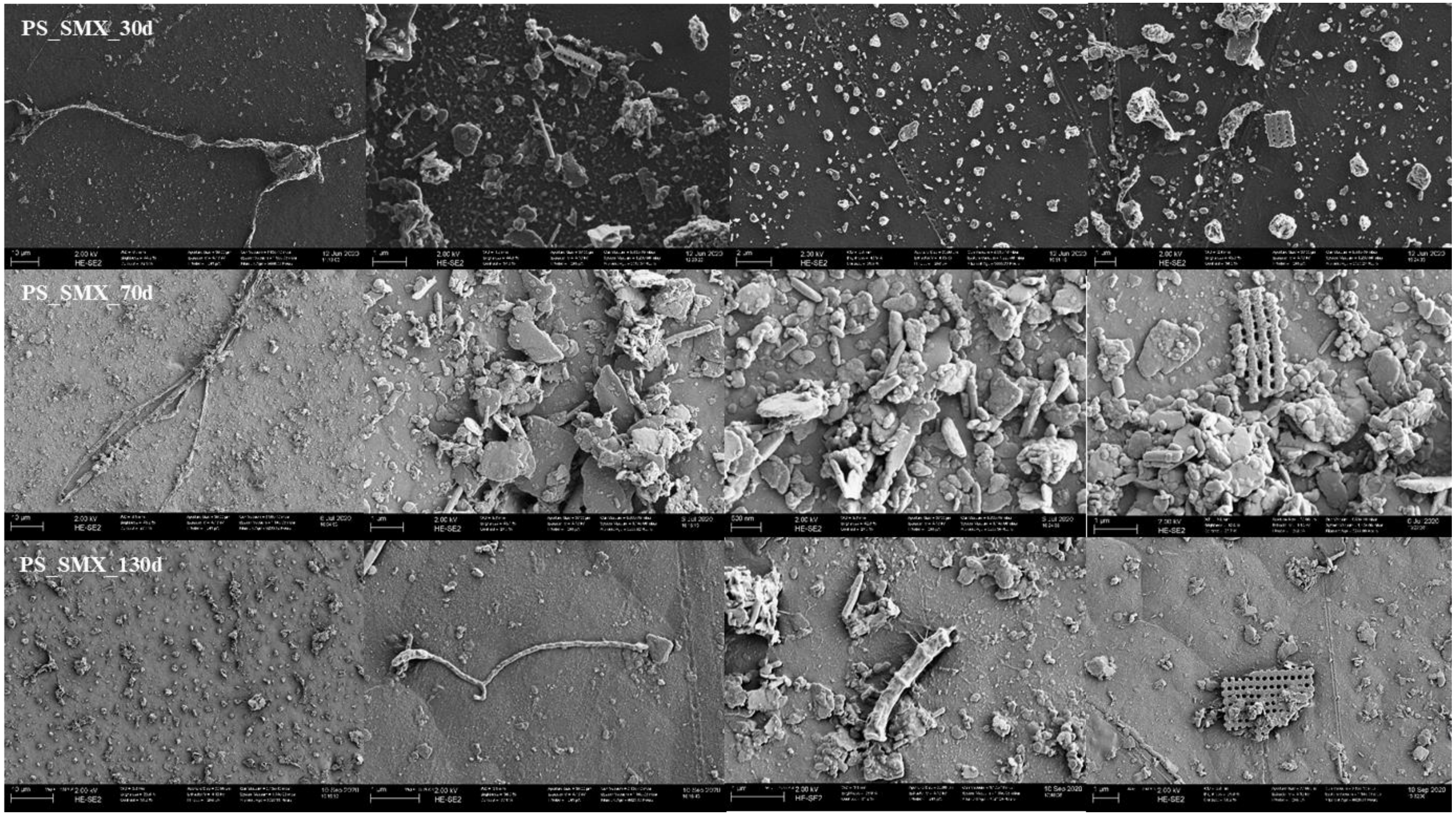


**Figure S10**. SEM images of microbial colonization of PS substrate incubated in soil for 30, 70 and 130 days with SMX amendment.

**Fig. S11**

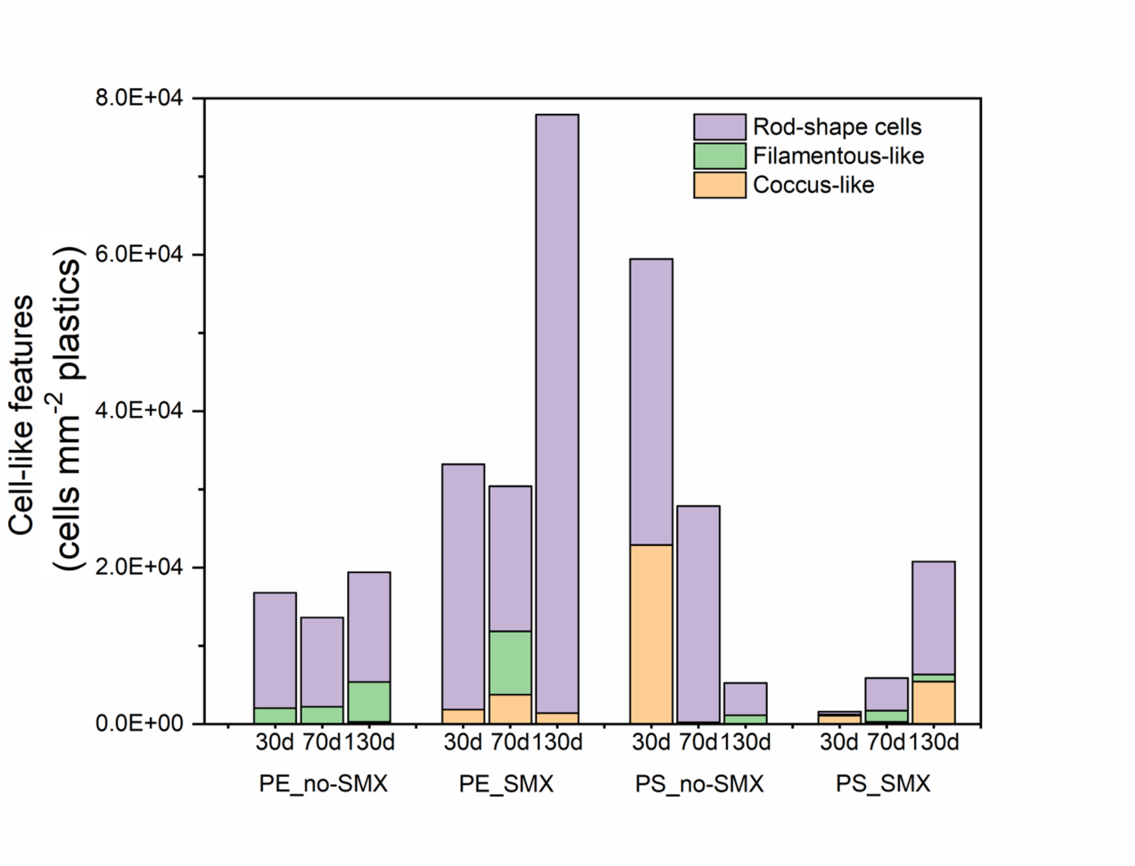


**Figure S11**. The stack bar charts showing the number of cell-like features observed in PE and PS plastisphere without or with SMX addition at different time points (30d, 70d and 130d).

**Fig. S12**

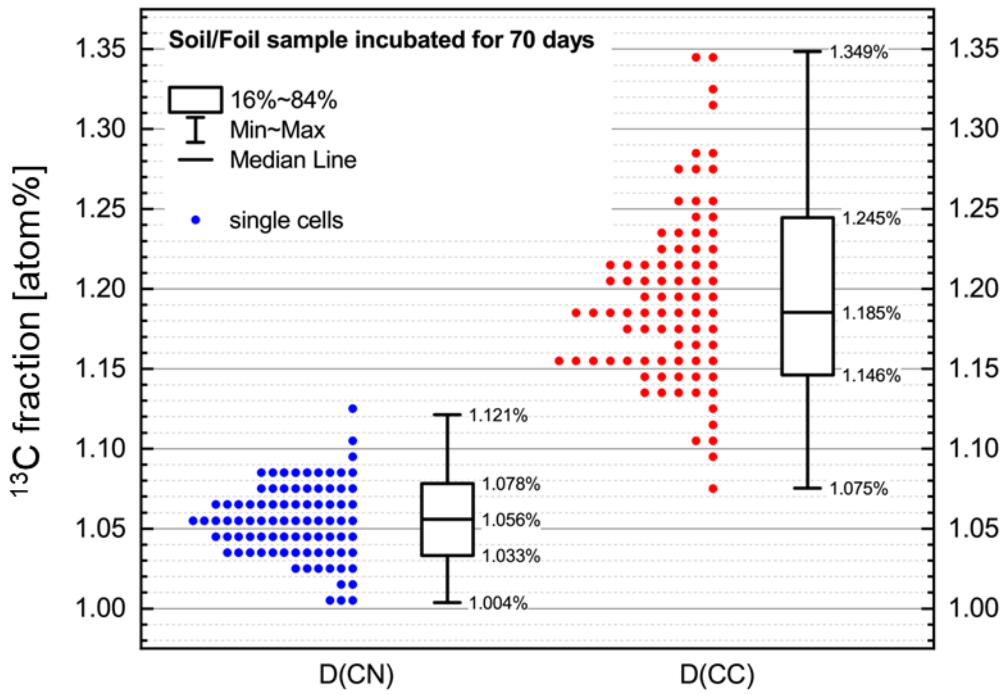


**Figure S12**. Distribution of the $^{13}C$ fraction derived from CN ($^{13}C^{14}N^-/^{12}C^{14}N^-$) and CC ($^{13}C_2{}^{12}C_2^-/^{12}C_2^-$) isotopic ratios of single cells colonizing the soil plastisphere PE at 70 day time point, without SMX.

**Table S1** The number of cells analyzed in the NanoSIMS single cell measurements.

<table>
<tr><th>Time / Plastic type</th><th>30d</th><th>70d</th><th>130d</th><th>Control</th></tr>
<tr><td>PE</td><td>103</td><td>84</td><td>103</td><td rowspan="2">85</td></tr>
<tr><td>PS</td><td>36</td><td>182</td><td>109</td></tr>
</table>
